# The speciation and hybridization history of the genus *Salmonella*

**DOI:** 10.1101/598235

**Authors:** Alexis Criscuolo, Sylvie Issenhuth-Jeanjean, Xavier Didelot, Kaisa Thorell, James Hale, Julian Parkhill, Nicholas R. Thomson, François-Xavier Weill, Daniel Falush, Sylvain Brisse

## Abstract

Bacteria and archaea make up most of natural diversity but the mechanisms that underlie the origin and maintenance of prokaryotic species are poorly understood. We investigated the speciation history of the genus *Salmonella*, an ecologically diverse bacterial lineage, within which *S. enterica* subsp. *enterica* is responsible for important human food-borne infections. We performed a survey of diversity across a large reference collection using multilocus sequence typing, followed by genome sequencing of distinct lineages. We identified eleven distinct phylogroups, three of which were previously undescribed. Strains assigned to *S. enterica* subsp. *salamae* are polyphyletic, with two distinct lineages that we designate Salamae A and Salamae B. Strains of subspecies *houtenae* are subdivided into two groups, Houtenae A and B and are both related to Selander’s group VII. A phylogroup we designate VIII was previously unknown. A simple binary fission model of speciation cannot explain observed patterns of sequence diversity. In the recent past, there have been large scale hybridization events involving an unsampled ancestral lineage and three distantly related lineages of the genus that have given rise to Houtenae A, Houtenae B and VII. We found no evidence for ongoing hybridization in the other eight lineages but detected more subtle signals of ancient recombination events. We are unable to fully resolve the speciation history of the genus, which might have involved additional speciation-by-hybridization or multi-way speciation events. Our results imply that traditional models of speciation by binary fission and divergence may not apply in *Salmonella*.

**Data summary:** Illumina sequence data were submitted to the European Nucleotide Archive under project number PRJEB2099 and are available from INSDC (NCBI/ENA/DDBJ) under accession numbers ERS011101 to ERS011146. The MLST sequence and profile data generated in this study have been publicly available on the *Salmonella* MLST web site between 2010 and the migration of the *Salmonella* MLST website to EnteroBase (https://enterobase.warwick.ac.uk/), and subsequently from there.

## Introduction

Bacteria and archaea make up most of natural diversity, both in terms of species richness and biological functions [1,2]. However, the mechanisms that underlie the origin and maintenance of prokaryotic species are poorly understood. It is often assumed that there is a single phylogenetic tree representing the relationships amongst prokaryotic taxa, with the branch lengths reflecting divergence times between them. However, bacteria and archaea acquire foreign DNA by homologous and non-homologous recombination and can recombine frequently, including in the *Salmonella* genus [3–9]. High recombination rates can maintain genetic cohesion within a species, preventing divergence and speciation from occurring until barriers to gene flow develop. Recombination has been shown in laboratory experiments to be supressed by nucleotide mismatches between donor and recipient [10,11]. This property provides a potential mechanism for speciation. It has been shown by simulation that large effective population sizes and neutral genetic drift can precipitate speciation by increasing the average pairwise divergence between strains, leading to either binary or multi-way speciation events [5,12].

Conversely, distinct new lineages or species can potentially arise almost instantaneously by hybridization of existing distantly related ones. Such large-scale hybridization events can occur at once by recombination of large genomic regions (e.g., [13]), or through multiple exchanges of small chromosomal segments associated with ecological convergence [14]. Therefore, to describe relationships between prokaryotes and understand patterns of species richness and phenotypic diversity, it is important to characterise the process of speciation and gene flow between species, including large-scale hybridization events [15].

Salmonellae are a prominent speciation model, where experimental and genomic studies of recombination and hybridization have been pioneered [4–10,14]. The genus *Salmonella* is divided into a number of phylogroups, namely *bongori, enterica, salamae, arizonae, diarizonae, houtenae*, and *indica* [16–18]. *Salmonella bongori* has been classified a distinct species [18], while the other phylogroups are considered to be subspecies of a single species, *S. enterica*. These taxa are further subdivided into serovars based on antigenic variation of flagellins and O-antigen.

Members of the genus *Salmonella* are major pathogens of humans and other warm-blooded animals. Human infections mostly involve *S. enterica* subspecies *enterica*, which can cause gastroenteritis, enteric fever and other infections [19,20]. Other *S. enterica* subspecies, as well as the species *S. bongori*, are more typically isolated from cold blooded animals or the environment, and are rarely reported from human infections [21].

Here we are concerned with evolutionary relationships rather than taxonomy and we designate phylogroups by names that derived from these subspecies’ labels, e.g. Bongori, Arizonae, Diarizonae, etc., with Enterica representing subspecies *enterica*. We use italicised names such as *houtenae* to refer to previous subspecies designations, which sometimes differ from our phylogroup assignments. A seventh *S. enterica* subgroup (group VII) was distinguished based on multilocus enzyme electrophoresis and gene sequencing [22–24]. Note that phylogenetic re-evaluation [25] of the proposed species *Salmonella subterranea* [26] shows that it does not belong to the *Salmonella* genus.

Phylogenetic analyses of the evolutionary relationships amongst the different *Salmonella* lineages have led to contradictory conclusions with several proposed phylogenetic trees [9,23,24,27–37]. This lack of consensus might reflect technical issues with phylogenetic reconstruction but a more biologically interesting possibility is that the history of *Salmonella* is not well-characterized by a simple model in which speciation proceeds stepwise by irreversible binary fissions.

To test this hypothesis, we sampled the genetic diversity within the little studied groups from cold-blooded hosts and used whole genome sequences from representative isolates of phylogroups to characterize the genetic relationships between them and to infer historical populations splits and gene flow. We show that while a binary fission model of speciation works for some of the *Salmonella* lineages, there are several important historical events that cannot be characterized in this way.

## Methods

### Taxonomic sampling and MLST analyses

A total of 367 strains (**Table S1**) from outside the subspecies enterica were selected from the collection of the World Health Organization Collaborative Centre for Reference and Research on *Salmonella*, Institut Pasteur, Paris, France. This center contains the reference strains of all *Salmonella* serovars and their variants. The 367 strains represented approximately one third of currently described serovars outside enterica and were selected to maximize the diversity of antigenic formulae. MLST was performed on these strains using updated primers adapted from those of Kidgell et al. [38] to amplify DNA from *S. bongori* and all subspecies of *S. enterica*. The novel primers are described in **Table S3**; note that they have been publicly available on the MLST web site between 2008 and the migration of the *Salmonella* MLST website to EnteroBase, and subsequently from there.

A phylogenetic tree was inferred from the median distance matrix of the seven genes with the algorithm BioNJ* [39]. A supermatrix of characters was built by concatenating the seven MSAs with the program Concatenate (www.supertriplets.univ-montp2.fr/PhyloTools.php), and the nucleotide diversity of groups was defined using the index π [40] with the program DnaSP [41]. Minimum spanning trees were built using the software tool BioNumerics (Applied-Maths, Belgium).

### Strain selection and genome sequencing

A set of 46 strains were selected for whole genome sequencing (**Table S2**). Genome sequencing was achieved by Illumina 2 x 50 nt paired-end sequencing for all strains. The characteristics of the obtained *de novo* assemblies are summarised in **Table S2**. This set was completed with genome sequences gathered from the GenBank repository (*i.e.*, 16 *S. enterica subsp. enterica*, 1 *S. enterica subsp. arizonae*, and 1 *S. bongori* strains), as well as 9 *S. bongori* genome sequences from Fookes et al. [34]. This led to a total of 73 genomes (**Table S2**): *S. enterica* subsp. *enterica*, 16; subsp. *salamae*, 13; subsp. *arizonae*, 9; subsp. *diarizonae*, 10; subsp. *houtenae*, 6; subsp. *indica*, 4; *S. bongori*, 10; and VII, 2.

### Core gene construction

Each of the 4,423 protein sequences from *S. enterica* strain LT2 [42] was used as query to perform BLAST similarity searches [43] against the genome sequence of each of the other 72 strains. Clusters of homologous sequences were built by considering only the first tblastn hit (E-value < 10^−5^), and every cluster that did not contain 73 sequences (*i.e.,* one per strain) was discarded. Next, orthology was assessed within each cluster by performing reciprocal tblastn, leading to 2,328 clusters of putative orthologous coding sequences from the core gene set of the 73 strains. For each of these clusters, sequences were translated, and a multiple sequence alignment (MSA) was performed with ProbCons [41] and next back-translated to obtain a codon-level MSA. The 2,328 MSAs were concatenated into a supermatrix of 2,137,446 nucleotide characters that was used to infer a balanced minimum-evolution phylogenetic tree using FastME [44] with pairwise p-distances. Branch support was assessed for each internal branch with a MSA-based bootstrap procedure based on 1,000 replicates. This procedure samples the MSA with replacement according to the same procedure as the standard bootstrap with nucleotide characters.

### Recombination analysis

We applied four separate and complementary methods to analyse the ancestral recombination events that occurred during the evolution of the genus *Salmonella*. Firstly, we applied chromosome painting on the 73 genomes, using CHROMOPAINTER [45] to reconstruct each genome as a mosaic of all the others. The results were summarized as a heatmap of coancestry values, where each coancestry value corresponds to the number of fragments copied from one genome to another (**Figure 2**). Secondly, we performed pairwise comparisons of genomes using a gene-by-gene approach. For each pair of genomes, we computed the genetic distances of all shared genes, and the distribution of these distances was plotted as a cumulative curve (**Figure 3**). Thirdly, the CHROMOPAINTER analysis was repeated using only nine unrelated genomes: one for each of the 12 phylogroups but excluding VII and Houtenae B due to recent shared ancestry with Houtenae A, and considering Enterica A and B as a single group. Each genome was therefore reconstructed as a mosaic of the other eight unrelated genomes. This allowed us to explore deeper relationships between phylogroups, since when all genomes are included each genome from a phylogroup copies mostly from other genomes of the same phylogroup (**Figure 2**). The resulting coancestry matrix was plotted as a heatmap (**Figure 4**). Fourthly, we applied the Treemix algorithm with parameter K=3 [46] to one genome from each of the 12 phylogroups in order to reconstruct their relationships as a vertical phylogenetic tree augmented with horizontal transfer events (**Figure 5**).

**Figure 1.**
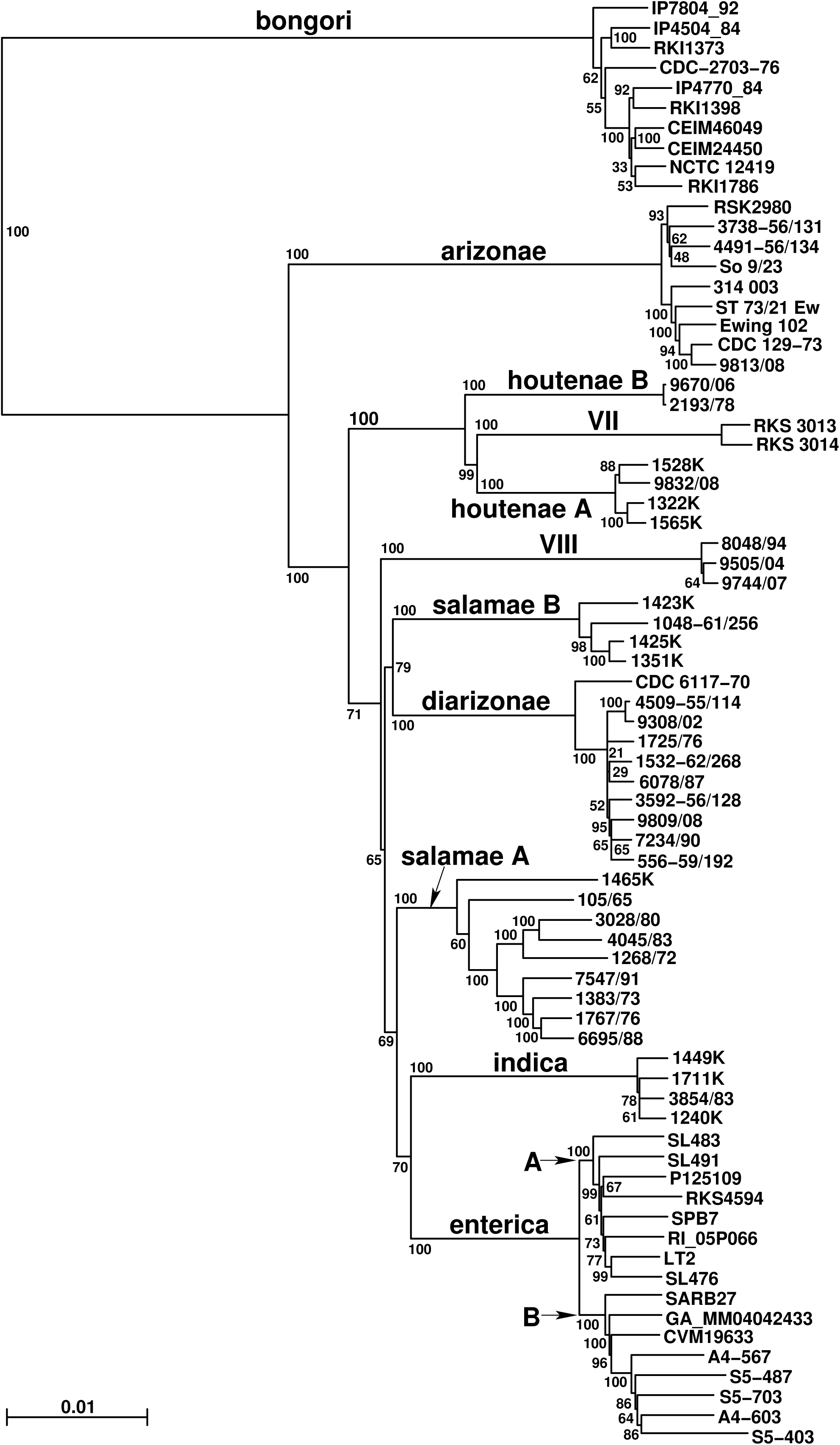
Phylogenetic tree of 73 *Salmonella* strains based on all shared core genes. The balanced minimum-evolution phylogenetic tree was constructed using FastME (see Methods). The 11 phylogroups are indicated above their ancestral branch; Enterica groups A and B are also indicated. Bootstrap/branch support values are indicated at the nodes.

**Figure 2.**
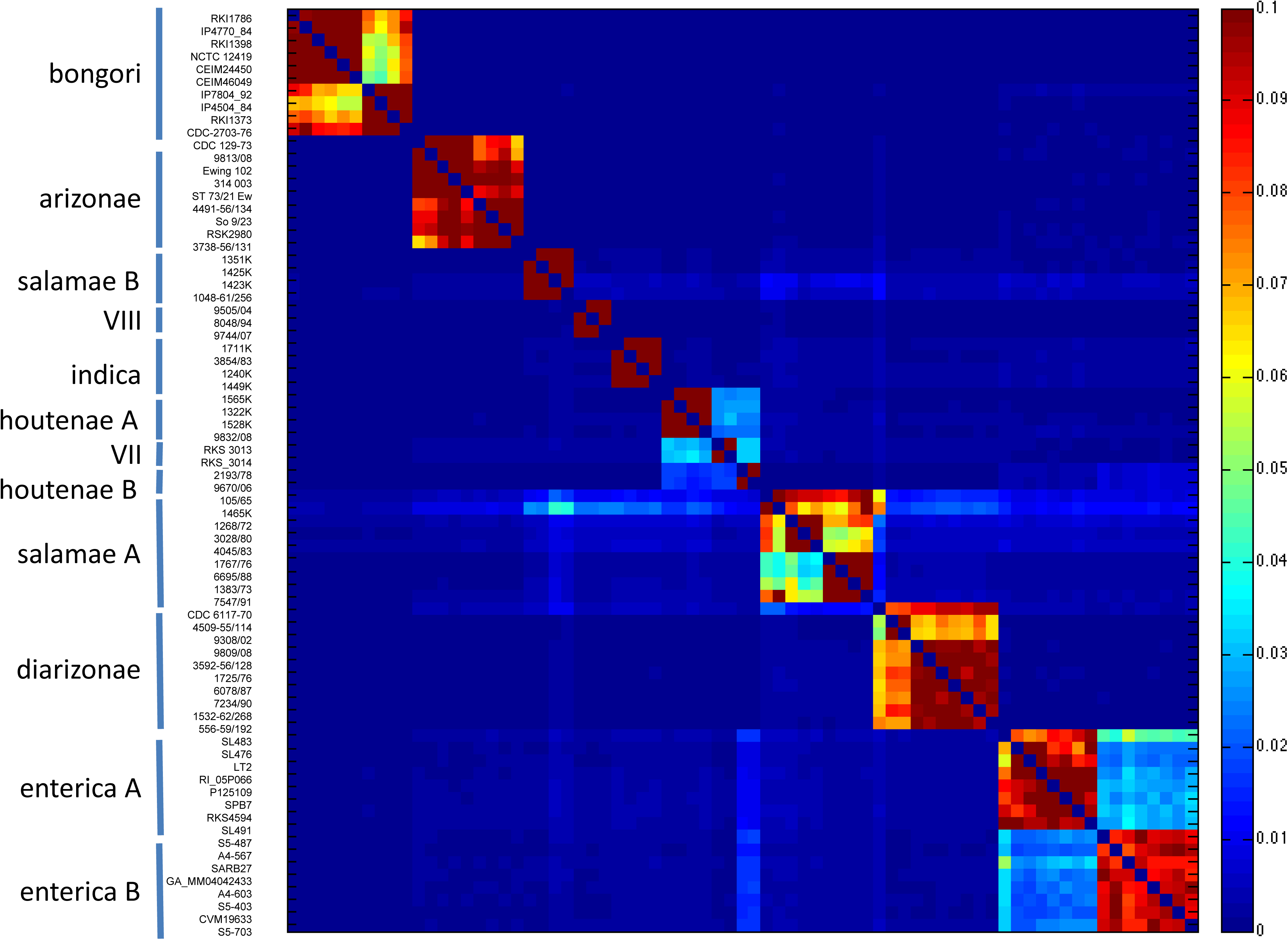
Coancestry matrix of 73 *Salmonella* genomes, computed using CHROMOPAINTER.

**Figure 3.**
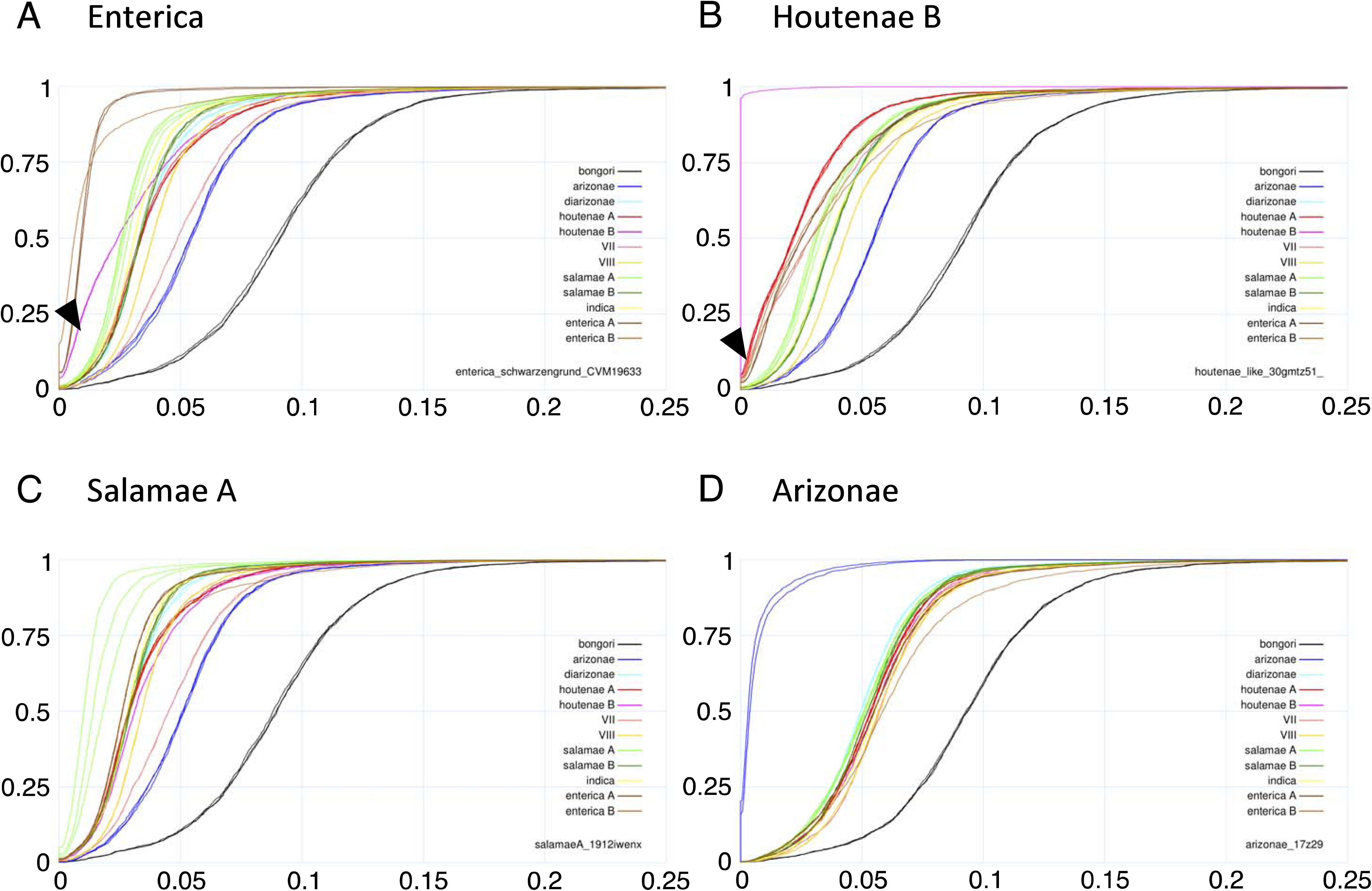
Cumulative curves of gene-by-gene distances between selected pairs of genomes. **A**: Comparisons with Enterica (group B, serovar Schwartzengrund CVM19633). The arrowhead shows that 20% (0.20, Y-axis) of the genes of an Enterica B strain have less than 1% (0.01, x-axis) divergence to Houtenae B. **B**: Comparisons with Houtenae B (2193/78). The arrowhead shows that 5% of the VII genome and 6% of Houtenae A has less than 0.1% divergence with Houtenae B. **C:** Comparisons with Salamae A (1268/72). **D:** Comparisons with Arizonae (CDC 129-73).

**Figure 4.**
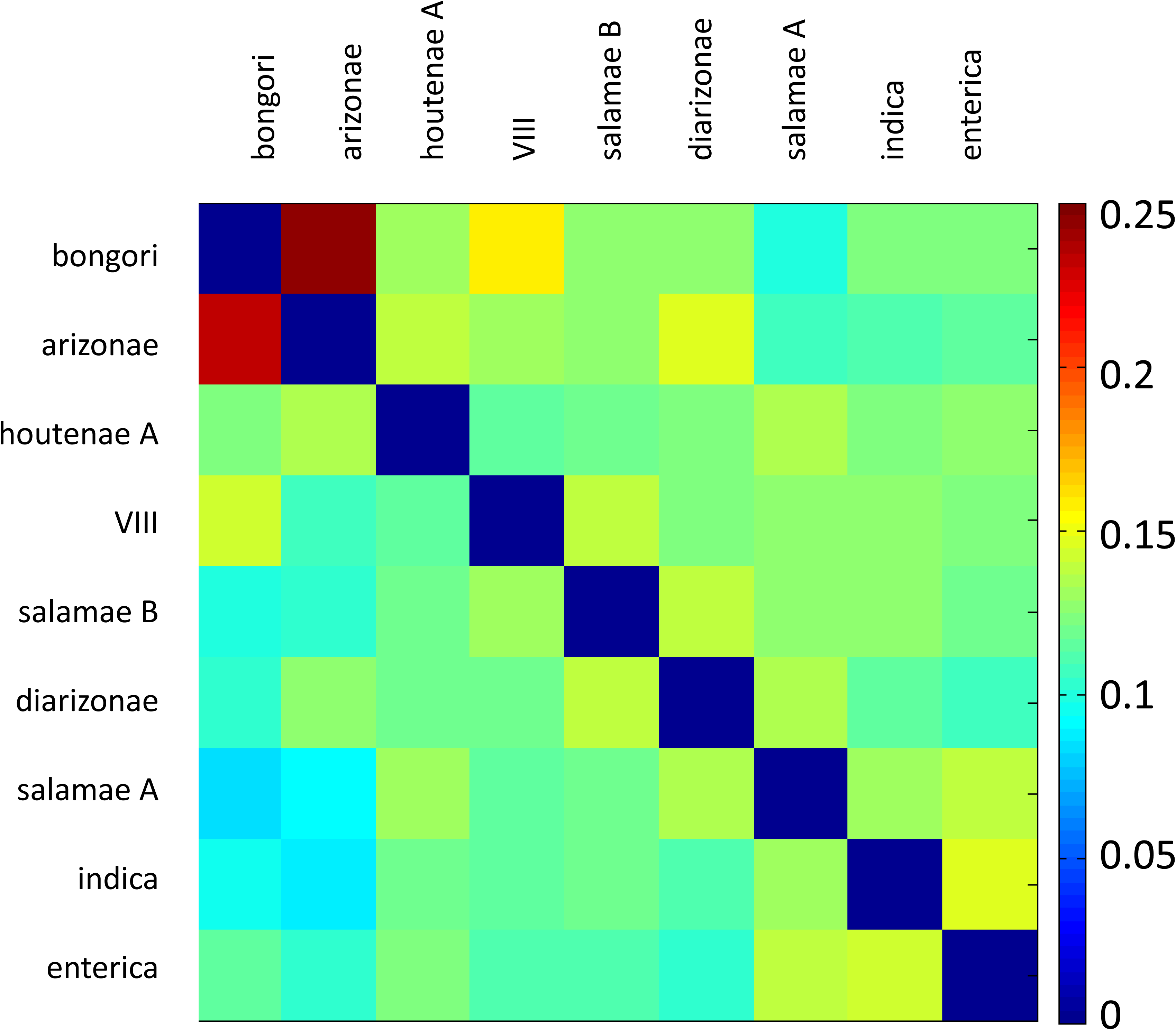
Coancestry matrix between 9 unrelated genomes, computed using CHROMOPAINTER.

**Figure 5.**
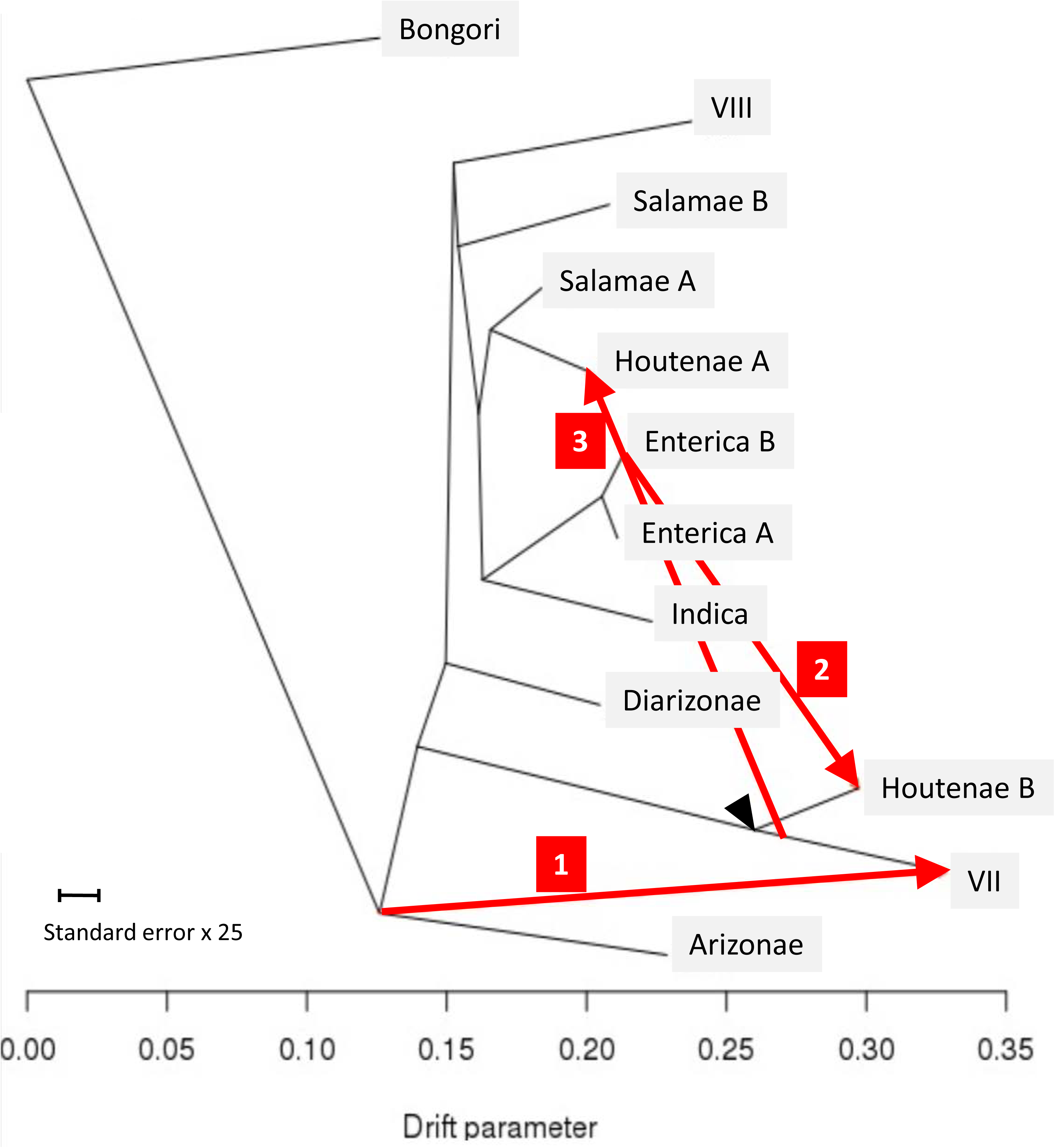
Treemix analysis of 12 genomes representative of phylogroups diversity. The arrowhead indicates the position of the ancestor contributing to extant Houtenae A, Houtenae B and VII lineages. The red arrows indicate gene fluxes inferred by Treemix.

### Pan-genome analyses

Analysis of accessory genome was performed using the Roary pan-genome pipeline version 3.6.2 [47]. Since the draft genomes were very unequally fragmented and synteny information therefore was of variable reliability we used the “don’t-split-paralogs” option. The analysis was performed with a protein identity cut-off of 85% and the core genome was defined as genes present in > 99% of the genomes studied. The Pearson correlation between accessory gene content of the genomes were visualised using the R software CORRPLOT package [48].

## Results and Discussion

In order to survey the diversity of *Salmonella* outside *S. enterica* subsp. *enterica*, a total of 367 strains, comprising about a third of the known non-Enterica serovars, were selected from the World Health Organization Collaborating Centre for Reference and Research on *Salmonella* (Institut Pasteur, Paris, France) reference collection and subjected to multilocus sequence typing (MLST) (**Tables S1, S2**). A phylogenetic tree was built (**Figure S1**), revealing a novel group (labelled VIII) and suggesting a polyphyletic origin of *S. enterica* subsp. *salamae* (Salamae A and B) and of *S. enterica* subsp. *houtenae* (Houtenae A and B). Within-phylogroup nucleotide diversity (**Figure S1 inset**) was the highest in Arizonae (π = 1.6%), lowest in Houtenae groups, Bongori, Salamae B and Diarizonae (π ranging from 0.35% to 0.42%), whereas it was intermediate in Salamae A, Indica and Enterica. Minimum spanning tree analysis of MLST profiles illustrates the genotypic diversity within each group (**Figure S2**).

Based on MLST diversity, 46 genomes were chosen for genome sequencing and compared to 27 previously published genome sequences of Enterica, Arizonae, and Bongori (**Table S2**). A phylogenetic tree based on the genome sequences is shown in **Figure 1**. This tree implies that *S. enterica* subsp. *salamae* is not a monophyletic group but instead forms two lineages with distinct evolutionary histories that we designate Salamae A and Salamae B. Whereas Salamae A contained 138 (88%) of the *salamae* strains, Salamae B comprised 18 isolates collected from a human (one isolate), a bat (one isolate) or reptiles (16 isolates, including 6 from chameleons). In contrast, 49 (41.5%) of Salamae A isolates were from humans, and only 34 (28.8%) were from cold-blooded animals, suggesting important ecological and pathogenic differences between the two Salamae groups. *S. enterica* subsp. *houtenae* was also subdivided into two distinct phylogroups, which we have designated Houtenae A and Houtenae B, and which clustered together with group VII on the tree. The genome-wide phylogenetic analysis also uncovers a hitherto unknown phylogroup, labelled VIII, made of strains previously identified as either salamae, diarizonae or of the former Hisingen serotype of *S. enterica* subsp. *enterica* [25]. The description of Salamae B, Houtenae B and VIII represent the first novel *Salmonella* phylogroups described since the identification of group VII by Selander and colleagues more than 25 years ago [24,27]. Our analysis therefore defines 11 phylogroups within *Salmonella*. The phylogenetic tree also shows further subdivisions at shallower levels, including the division of *S. enterica* subsp. *enterica* into Enterica A and Enterica B as previously described [5,9]. Note that the genomes of the present study have been publicly available from International Nucleotide Sequence Database Collaboration (INSDC) since 2011, and were used in a genome-based phylogenetic analysis of *Salmonella* by Alikhan *et al*. [49]; the three novel *Salmonella* groups were labelled as novel subspecies A (Houtenae B), B (VIII) and C (Salamae B) in [49].

### Recent recombination between phylogroups

We used chromosome painting of the above set of 73 strains in order to investigate shared ancestry and recombination events between different phylogroups. Specifically, the CHROMOPAINTER algorithm uses a Hidden Markov Model to reconstruct each isolate as a mosaic of stretches of DNA of the other isolates in the sample [45]. The results can be summarized as a heatmap indicating how many stretches from each other sample are used in the reconstruction. The organism used in the reconstruction is assumed to be the most closely related for each stretch of DNA. **Figure 2** shows a heatmap illustrating the proportion of DNA used to paint each isolate across the genome, with dark blue corresponding to 0% and dark red corresponding to 10%. We call this proportion the coancestry value. Each phylogroup shows higher coancestry within the same phylogroup than with others. The highest coancestry between strains in different phylogroups is between Houtenae A, Houtenae B and VII. However, Houtenae B shows higher Enterica ancestry (particularly with Enterica B) than do Houtenae A or VII. The two deepest branching Salamae A strains show high levels of coancestry with several other groups including Salamae B, Diarizonae, Indica and VIII. One strain of Enterica A (SL483) is exceptional in showing higher coancestry levels with Enterica B.

In order to test whether high coancestry between groups might be explained by recent recombination between them, we looked for evidence of sharing of very similar stretches of DNA between pairs of lineages [14] by plotting, for each pairwise comparison, the proportion of genes with divergence below a threshold increasing from 0% to 25% (**Figure 3**). Consistent with recent recombination between them, Enterica B and Houtenae B showed many more genes with very similar sequences than expected based on their position in the phylogenetic tree, with 20% of the genes of an Enterica B strain having less than 1% divergence to Houtenae B, compared to only 5% between Enterica B and Houtenae A (**Figure 3A**). These divergence curves are also consistent with recent recombination between Houtenae A, Houtenae B and VII. For example, approximately 5% of the VII genome and 6% of Houtenae A has less than 0.1% divergence with Houtenae B (**Figure 3B**), suggesting that there has been very recent recombination between these three phylogroups. There is no analogous signal of recent recombination between any of the strains of Salamae A or Salamae B with each other or with other phylogroups based on cumulative divergence curves (*e.g*., **Figure 3C**). The smudged pattern of coancestry of the deeper branching Salamae A and Salamae B strains in **Figure 2** can potentially be explained by them retaining ancestral variants that have been lost by the rest of the phylogroup and therefore does not necessarily indicate recent recombination between lineages. **Figure 3D** illustrates the absence of any signal of recent recombination with Arizonae.

### Evidence for hybridization in the origin of the phylogroups

We next examined the origins of the phylogroups themselves. Recombination events which predate the generation of the diversity observed *within* each phylogroup are unlikely to be picked up in the chromosome painting analysis in **Figure 2**: members of a phylogroup that have inherited the same ancestrally imported stretch will be painted by each other for those stretches. Therefore, we selected a single strain from each phylogroup and performed a distinct chromosome painting analysis. We excluded VII and Houtenae B due to the recent shared ancestry with Houtenae A, and also included only a single representative for both Enterica A and Enterica B. The chromosome painting results (**Figure 4**) show high coancestry between Bongori and Arizonae and between Indica and Enterica. These relationships can be interpreted using a vertical phylogenetic model, as they agree with a large number of different analyses including ours (**Figure 1)** that Arizonae is the deepest branching lineage after Bongori and that Indica is a sister group of Enterica [9,24,33,34]. Conversely, the chromosome painting analysis reveals a large number of intransitive relationships (*i.e*., in which A has elevated coancestry with B and B has high coancestry with C but C does not have high coancestry with A). First, Diarizonae and Arizonae have high coancestry, as do Diarizonae and Salamae B but Salamae B and Arizonae do not (**Figure 4**). Second, Houtenae A and Salamae A have high coancestry with each other and the phylogenetic tree suggests they are sister taxa. However, they have different relationships to other phylogroups. Houtenae A, but not Salamae A, shows high coancestry with Arizonae, while Salamae A shows higher shared ancestry with Indica and Enterica. Intransitive patterns of coancestry are also evident for VIII, Salamae B and Diarizonae and for VIII, Salamae B and Bongori. An intransitive pattern is not predicted by any phylogenetic model and is indicative of mixture in the history. These observations suggest a complex pattern of homologous recombination events that predate diversification within phylogroups.

### A scenario involving three recent hybridization events

To complement the results above, we used Treemix to infer a history that allows for recombination events in the origins of the phylogroups. Treemix attempts to model the covariance matrix reflecting SNP sharing between strains by assuming a phylogenetic model of divergence via genetic drift, but with a limited number *K* of mixing events in the history. Our application of Treemix to *Salmonella* gave results which varied in important details depending on the value of K. Each of the events that were identified at a given value of K had counterparts in the inference performed for higher values, but details of the inferred phylogenetic tree and the location and direction of the hybridization events were not consistent. For example, for K=1 and K=2 Houtenae A and Houtenae B are sister taxa whose common ancestor received genetic material from VII, while for K=3, VII and Houtenae B share a common ancestor, which contributed genetic material to Houtenae A. We present the Treemix results for K=3 (**Figure 5**) because all of the events inferred are supported by signals identified by chromosome painting and cumulative divergence (**Figures 2, 3 and 4**). The Treemix results with K=3 imply that Houtenae A, Houtenae B and VII all have hybrid origins. All three of them received DNA from a shared lineage which branched between Arizonae and Diarizonae (black arrowhead, **Figure 5**), but differ in the remaining source of their ancestry (red arrows, **Figure 5**), which, according to the Treemix estimates, account for about half of their genome in all three cases (1: ancestor of Arizonae to VII: 0.461; 2: Enterica B to Houtenae B: 0.42; 3: ancestor of VII to houtenae A: 0.49). Note that according to this reconstruction, no pure, or nearly pure, representative of this shared ancestral lineage is present in the sample, a feature which is likely to have contributed largely to the instability of the Treemix analysis and makes all types of evolutionary reconstruction considerably more challenging.

The second source for Houtenae B is inferred to be Enterica B (red arrow 2, **Figure 5**), which is consistent with the results from chromosome painting and of the pairwise distances, as discussed above, and is consistent with recent genetic exchange having taken place. The second source for VII is inferred to branch at the same point as Arizonae does in the tree. The deep position of this ancestry source is supported by the distribution of pairwise distances VII has to shallower branching lineages such as Diarizonae or Salamae A, which are more similar to the distribution found for Arizonae than to that of either Houtenae A or Houtenae B (*e.g*., **Figure 3C**). The distribution of distances to Arizonae is similar to that of other shallow-branching lineages, suggesting that the recombination was not with Arizonae itself. Finally, the second source for Houtenae A branches next to Salamae A, which is consistent with the reconstructed position of Houtenae A in the phylogenetic tree in Figure 1 and the high coancestry of Houtenae A and Salamae A in Figure 4. However, unlike for Houtenae B, there is no signal of recent recombination of Houtenae A with other lineages in Figure 2. Furthermore, the pairwise distance curves of Salamae A to Houtenae A and Houtenae B are comparable (**Figure 3C**). These features imply that there has not been recent recombination between Houtenae A and Salamae A. Instead, they are consistent with the second source that contributed to Houtenae A being an unsampled sister taxa to Salamae A.

### Unequal evolutionary rates of the different taxa

One important feature of the phylogenetic tree (**Figure 1**) is the different branch lengths leading to each phylogroup. This feature might be caused by either unequal substitution rates between lineages or by recombination, which can cause hybrid lineages to branch closer to the root. Evidence for unequal substitution rates comes for example from comparisons with Bongori or Arizonae, which can tentatively be treated as outgroups. Salamae A and Salamae B have smaller genetic distances than other lineages to either (**Figure 3D**), despite the chromosome painting indicating no evidence of elevated recombination between them. Furthermore, Salamae A and Salamae B show low genetic distances compared to potential sister lineages to all taxa, suggesting that they have substantially lower substitution rates than other groups. Because our reconstruction of *Salmonella* evolutionary history is incomplete and uncertain, we do not attempt to formally model all of these processes occurring together.

### Accessory genome relationships

Accessory genes contribute most to ecological specialization and the pattern of horizontal gene transfer among phylogroups might provide important complementary information regarding functional and ecological correlates of the recombination history that we inferred in this work [9]. We therefore analysed the pan-genome of the dataset, which with a protein identity cut-off of 85% rendered a core genome of 1818 gene clusters and a total pan-genome of 21973 genes. Unfortunately, estimations of the strain relationships based on gene presence/absence and analysis of the shared ancestry revealed that the analyses were strongly affected by the fragmentation of the genomic assemblies (**Table S2**), as was particularly visible for the highly fragmented Diarizonae genomes (**Figure S3)**. Analysis of the horizontal gene transfer pattern among phylogroups therefore requires higher quality assemblies and will be the subject of future studies.

## Conclusions

We investigated the diversification and hybridization history within *Salmonella*, a group of prominent public health importance and an early model for microbial speciation and evolutionary studies. By sampling largely in the non-enterica subspecies, we uncovered three novel phylogenetic groups that had not been recognized since the last group, VII, was described in 1991. Our snapshot of diversity within phylogroups of *Salmonella* implies that recombination among phylogroups is relatively rare at any point in time but that when it happens it can be with distantly related lineages rather than sister taxa and can involve large fractions of the core genome. These events are likely to provide substantial potential for phenotypic innovation but may also entail a great deal of hybrid disgenesis.

The three hybridization events that we have been able to elucidate with any degree of certainty are ongoing or took place in the recent past and all involved a lineage that is not present in unhybridized form in the dataset. This circumstance makes it challenging to estimate simple properties of the events such as the direction of hybridization and the proportion of genome acquired from each source. We can nevertheless robustly conclude that the hybridization has involved at least three entirely different branches of the *Salmonella* tree and has led to the formation of three phylogroups, namely Houtenae A, Houtenae B and VII. Interestingly the latter group was inferred to be a ‘hybrid’ in early MLEE studies [24]. This suggests an interesting question that is likely to be informative about the general nature of species boundaries in bacteria, namely what has happened to make one lineage particularly prone to hybridization in the recent past?

We see less conclusive but nevertheless still strong evidence for hybridization events in the more distant past. Phylogenetic trees of *Salmonella* phylogroups are notoriously unstable, including in different analyses we have performed (data not shown). In particular, relationships amongst Salamae A, Salamae B, Diarizonae, Enterica and VIII are difficult to elucidate. The coancestry relationships between these lineages are highly intransitive (**Figure 4**). One possibility is that this intransitivity is due to a complex multi-way speciation event [5], such that there is no true splitting order to infer. However, it may also represent hybridization events after stepwise speciation. The two lineages that branch deeply (**Figure 1**), namely VIII and Diarizonae, both show evidence of shared ancestry with basal lineages, Bongori and Arizonae, respectively (**Figure 4**), which is likely to have substantially affected their phylogenetic position.

The events of recombination inferred in this work explain the difficulties to reconstruct the phylogeny of the genus that have led to multiple distinct hypotheses on the phylogenetic relationships among subspecies. The phylogenetic relationships which do appear to be reasonably certain are that Bongori split from the other phylogroups first, followed by Arizonae and that Indica is a sister group to Enterica. Houtenae A seems to have been a sister taxon of Salamae A, prior to its mixture event. These examples demonstrate that in the right circumstances, phylogenetic signal can be preserved over long evolutionary time periods despite recombination between phylogroups. The problem of reconstructing ancestral hybridization events is a hard one and we do not have the tools or genomes available to reconstruct an entire history with any degree of confidence.

Our results demonstrate that bacterial species histories are complex. There is considerable phylogenetic signal in the data, consistent with the evolution and long-term persistence barriers to gene flow between lineages but also examples for hybridization events that may reverse species boundaries, sometimes between taxa separated by large genetic distances, rather than between sister taxa. These results mean that phylogenetic trees displaying relationships between species will often represent considerable simplifications of evolutionary history and in the worst case can be entirely misleading. Further work in multiple taxa will elucidate the evolutionary and ecological factors that precipitate speciation and hybridization events.

## Author contributions

***Conceptualization: SB, DF. Supervision:* SB, DF, NRT, FXW. *Performed the experiments:* SIJ. *Data curation:* JH, SB, SIJ, FXW. *Data analysis:* AC, SB, XD, KT. *Writing – original draft:* AC, SB, DF. *Writing – review & editing:* all.**

## Acknowledgements

We acknowledge the expert help of Laure Diancourt, Coralie Tran and Virginie Passet (Institut Pasteur) for MLST data production, and of Mark Achtman for his support at the start of the project and for bioinformatics assistance in MLST data curation.

## Supporting Information

**Table S1. Strains studied by MLST.**

**Table S2. Genomic sequence data.**

**Table S3. Primers and conditions used for MLST gene amplification and sequencing for non-Enterica isolates.**

**Figure S1.**
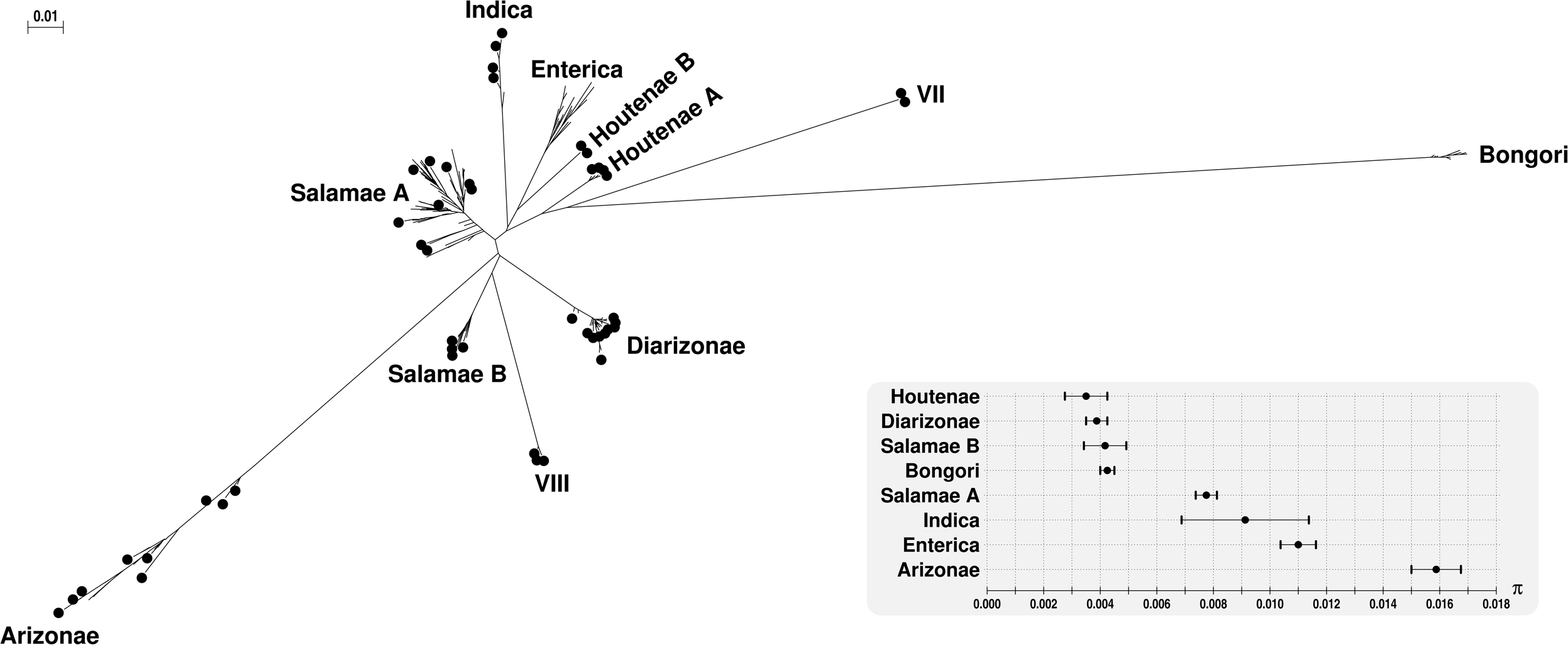
BioNJ* tree of 382 *Salmonella* strains based on seven housekeeping gene sequences. The inset shows the average nucleotide diversity of each phylogroup (houtenae comprises Houtenae A and Houtenae B) at the seven MLST genes.

**Figure S2.**
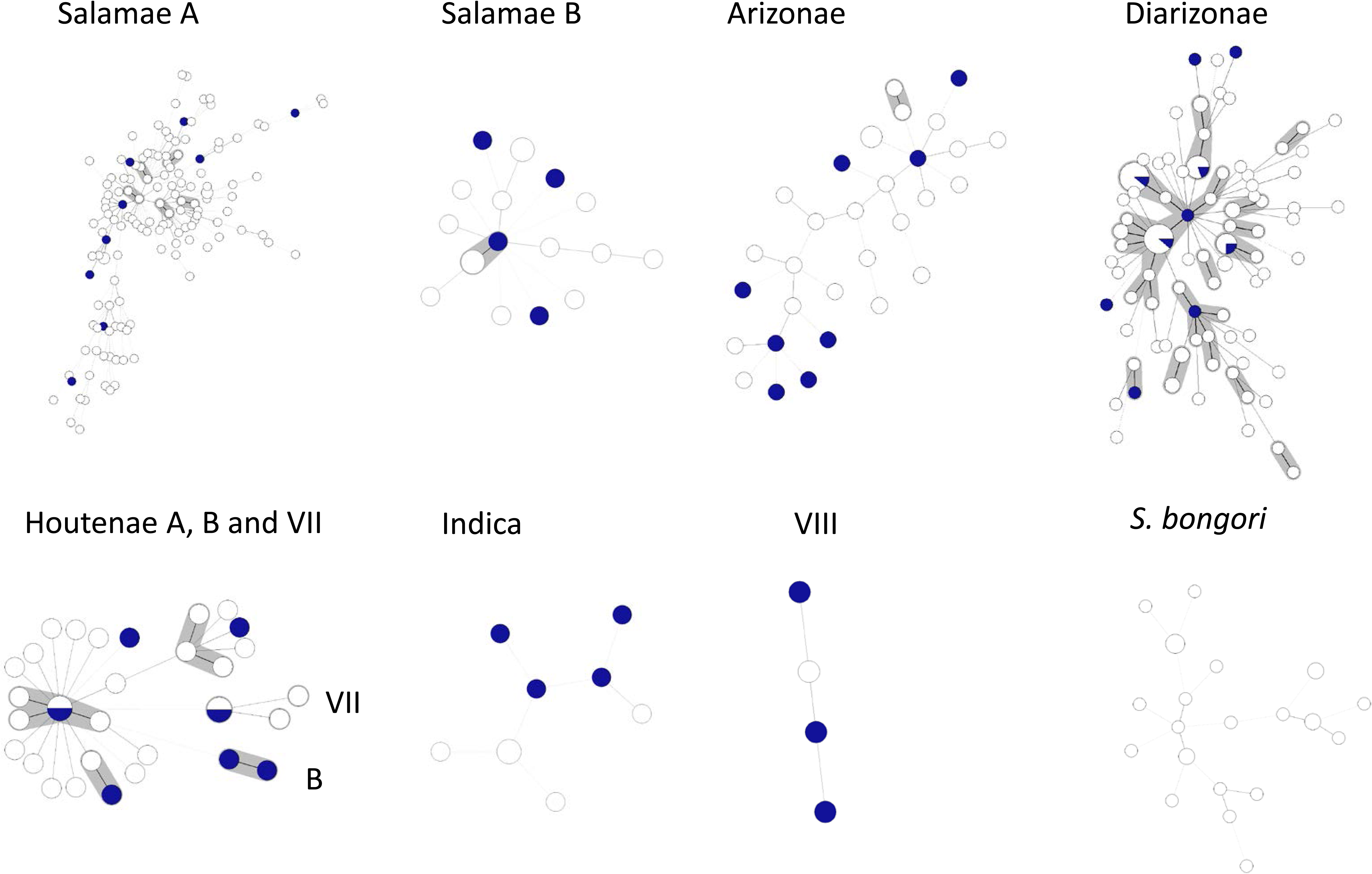
Minimum spanning tree representations of the genotypic diversity within *Salmonella* groups. The minimum spanning trees were constructed for each group based on number of mismatches among MLST allelic profiles. Strains selected for genome sequencing are represented by blue sectors (or blue circles when only one strain shared the corresponding genotype). Grey zones surround groups of sequence types that are connected successively by single allelic mismatches and are equivalent to clonal complexes or ‘eBURST’ groups (Achtman et al., 2012).

**Figure S3.**
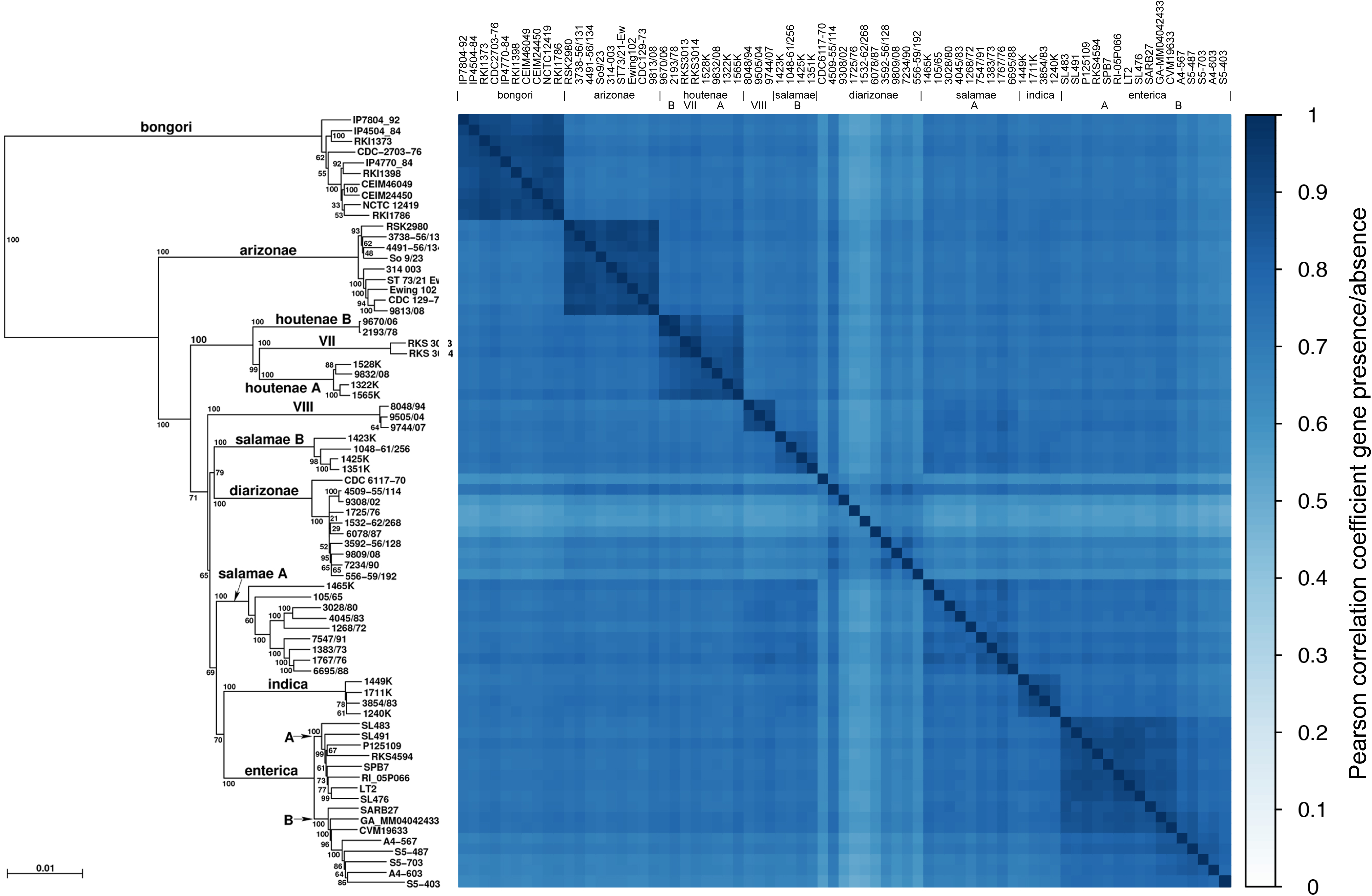
Heatmap of the proportion of shared genes. Strains are ordered according to the phylogeny in Figure 1 (left). The proportion of shared genes was computed from the ROARY output with a protein identity cut-off of 85% and the “don’t split paralogs” option.

